# Genome-wide association mapping of transcriptome variation in *Mimulus guttatus* indicates differing patterns of selection on cis- versus trans-acting mutations

**DOI:** 10.1101/2021.06.02.446804

**Authors:** Keely E. Brown, John K. Kelly

## Abstract

We measured the floral bud transcriptome of 151 fully sequenced lines of *Mimulus guttatus* from one natural population. Thousands of single nucleotide polymorphisms (SNPs) are implicated as transcription regulators, but there is a striking difference in the Allele Frequency Spectrum (AFS) of cis-acting and trans-acting mutations. Cis-SNPs have intermediate frequencies (consistent with balancing selection) while trans-SNPs exhibit a rare-alleles model (consistent with purifying selection). This pattern only becomes clear when transcript variation is normalized on a gene-to-gene basis. If a global normalization is applied, as is typically in RNAseq experiments, asymmetric transcript distributions combined with “rarity disequilibrium” produce a super-abundance of false positives for trans-acting SNPs. To explore the cause of purifying selection on trans-acting mutations, we identified gene expression modules as sets of co-expressed genes. The extent to which trans-acting mutations influence modules is a strong predictor of allele frequency. Mutations altering expression of genes with high “connectedness” (those that are highly predictive of the representative module expression value) have the lowest allele frequency. The expression modules can also predict whole-plant traits such as flower size. We find that a substantial portion of the genetic (co)variance among traits can be described as an emergent property of genetic effects on expression modules.

---

Genetically controlled gene expression variation is prevalent within and between species and across different tissues, environments, and treatment contexts (Harding et al. 1989, Whitehead and Crawford 2006, McManus et al. 2010, Meiklejohn et al. 2014, Signor and Nuzhdin 2018). Changes in gene expression can facilitate divergence between species (Johnson and Porter 2000, Tulchinsky et al. 2014, Mack and Nachman 2017, McGirr and Martin 2020), and provide a mechanism for a population to rapidly adapt to a new environment (Morris et al. 2014, Ghalambor et al. 2015, Campbell-Staton et al. 2017, Margres et al. 2017, Mack et al. 2018, Hamann et al. 2020). Standing genetic variation and plasticity in gene expression can buffer a population against environmental fluctuations (Podrabsky and Somero 2004, Stern et al. 2007, Acar et al. 2008, López-Maury et al. 2008). While much has been learned about the regulation of particular genes, genome-wide patterns in the evolutionary dynamics of gene expression are just beginning to be explored (e.g. Josephs et al. 2020). It also remains unclear how gene expression, as a molecular phenotype, might mediate the genetic underpinnings of quantitative trait variation, and ultimately fitness.

## Evolutionary dynamics of transcriptional effectors

Mutations can alter gene expression in many ways, and we expect selection to act differently on different types of variants (Lawrence et al. 2016, Bewick and Schmitz 2017, Duren et al. 2017). Gene expression can be affected by mutations acting either in cis or in trans. Cis-acting variants affect a closely linked gene directly, perhaps by altering sequences normally bound by transcription factors or other regulatory machinery. Trans-acting regulatory variants, in contrast, change the cellular environment in which transcription happens, say by altering diffusible products like the transcription factors themselves (Wittkopp et al 2004, Emerson and Li 2010).

Natural selection could differ systematically between cis- and trans-acting variants for several reasons. First, the mutational target for trans-acting effectors of a gene could be a substantial fraction of the genome (Boyle et al. 2017), compared to the more limited set of sites at which cis-acting mutations could occur (Gruber et al. 2012, Metzger et al. 2016). Second, trans-acting variants have the potential to affect multiple genes, and are may thus have negative consequences on finely-tuned pathways (Stern and Orgogozo 2008; but see Hoekstra and Coyne, 2007). If trans-mutations have opposing pleiotropic effects on many genes (antagonism), they may still increase in frequency in a conditional manner (Hall et al. 2010, Anderson et al. 2011). Third, if a trans-acting variant with weakly deleterious effects on the expression of one or more target genes increases substantially in frequency (due to drift or selection), cis-variants specific to each affected gene might then act as a buffer, leading to positive directional selection on cis-compensatory mutations. This often occurs, for example, with pleiotropic mutations associated with antibiotic resistance (Maisnier-Patin and Andersson 2004, Brandis et al. 2012) and compensatory pairs of cis- and trans-effectors have been documented in several systems (Coolon et al. 2014, Wang et al. 2015, Fear et al. 2016, Mack et al. 2016, Verta et al. 2016, Metzger et al. 2017). These theories generally suggest that trans-acting variants should be under stronger negative selection than cis-acting variants based on the premise that gene expression should usually experience strong stabilizing selection (Denver et al. 2005, Rifkin et al. 2005, Whitehead and Crawford 2006, Hodgins-Davis et al. 2015). If this is correct, then any mutation with broad effects on expression, regardless of cis- or trans-effect, will more likely be deleterious, perhaps through cascading effects on connected pathways or networks (Fisher 1930).

Broad patterns of selection can be inferred from the allele frequency spectrum (AFS) of cis- and trans-acting variants. When compared to the neutral expectation, an excess of intermediate frequency variants suggests balancing selection while an excess of rare variants suggests purifying selection (Tajima, 1989). Demographic events, such as population expansions or contractions, can perturb the AFS away from the neutral expectation (Hartl and Clark, 1997). However, since these effects are genome-wide, we can make inferences about selection by comparing the AFS for a particular class of polymorphism (e.g. cis-effectors of gene expression) to that of the entire genome. Of course, this is just a first step; inferences about selection require corroboration from multiple lines of evidence (Beaumont and Balding 2004, Bigham et al. 2010). In this study, we find an AFS consistent with purifying selection for trans-acting expression variants and corroborate this pattern by showing that the skew towards extreme allele frequencies is greatest at loci with the broadest effects on expression. In contrast to trans-SNPs, cis-regulators exhibit an AFS suggesting balancing selection where corroboration is more difficult. The processes most likely to generate balancing selection on cis-SNPs depend on the specific ways that these mutations affect whole organism phenotypes, and also on the complicated and variable mapping from phenotype to fitness in nature.

## Transcriptional mutations generating genetic covariation in traits

How important are transcriptional regulators in modifying fitness-related traits? Case studies of specific genes with known mutant phenotypes provide many examples where gene expression influences fitness relevant traits of plants (Streisfeld and Rausher 2009, Sobel and Streisfeld 2013, Ning et al. 2017, Kremling et al. 2018, Alonge et al. 2020). The quantitative importance of transcriptional mutations relative to those that effect enzymatic or structural protein function remains a point of contention (Hoekstra and Coyne 2007, Stern and Orgogozo 2008), but a steady increase of evidence from human eQTL/eGWAS research suggests a predominant role for gene expression variation in generating quantitative trait variation (Nicolae et al. 2010, Maurano et al. 2012, Torres et al. 2014, Farh et al. 2015, Boyle et al. 2017). As a first step to understanding the selection regime on cis-acting variants, we here use observed gene expression variation to predict variation and co-variation among a set of quantitative traits of yellow monkeyflower (*Mimulus guttatus)*. These traits were previously analyzed as part of a GWAS that predicted trait values and fitness estimates in nature directly from SNPs (Troth et al 2018).

Understanding trait covariances is as important as trait variances when natural selection involves trade-offs between different traits. Such trade-offs can provide the mechanistic basis of balancing selection (Mérot et al. 2020). In many annual plants, suites of correlated life-history traits related to rate of development (progression to flowering) are subject to a tradeoff between flowering time and fecundity. In Mimulus specifically, variation in life-history traits is maintained by opposing selective pressures on survival to flower and seed set (Kelly 2008, Mojica and Kelly 2010, Mojica et al. 2012, Monnahan and Kelly 2015, Troth et al. 2018). As with many other species, small, fast-growing plants survive to flower but make fewer seeds. Large, slow-growing plants make more seeds but risk not reaching maturity before the end of the growing season. This type of tradeoff can maintain polymorphism at loci affecting the underlying traits such as days to flower or flower size (Austen et al. 2017, Brown and Kelly 2018, Exposito-Alonso et al. 2018). While previous work has mapped this trade-off to the pleiotropic effects of specific genetic loci, it remains unclear if gene expression mediates these effects.

Here, we associate SNPs segregating within inbred lines derived from the Iron Mountain population with gene expression variation in flower buds (an “eGWAS” or “eQTL” study). Allele frequencies in the inbred lines accurately represent those in the natural population (Troth et al. 2018). We first document strikingly different patterns of apparent selection from the AFS of cis- and trans-acting regulatory SNPs. We then show that modules of co-expressed genes predict the trait means of the inbred lines, despite that we measured gene expression and traits on different plants grown in different places. The stability of the relationship between transcriptome and trait is surprising, given the supposed “noisiness” of transcriptome data (Arias and Hayward 2006, Raj and Oudenaarden 2008). Finally, we demonstrate that correlations, including tradeoffs between fitness-related traits can be predicted from gene expression variation.

## Methods

### Study system

We used randomly derived inbred lines of the yellow monkeyflower, *Mimulus guttatus (*syn *Erythranthe guttata,* Phrymaceae) from the Iron Mountain (IM) population in the Cascade Mountains of Oregon (44.402217N, 122.153317W) (Willis 1999, Kelly 2003). This population is predominantly outcrossing with little internal population structure (Sweigart et al. 1999, Willis 1993). Due to its annual/winter annual lifespan and short growing season, the IM population experiences a fitness tradeoff caused by variation in flower size and life-history phenotypes (Mojica et al. 2012). In 2018, Troth et al. sequenced whole genomes of 187 IM inbred lines and phenotyped them for 13 flower size and developmental timing traits known to influence fitness in the field.

### RNAseq

We grew plants from 151 of the genome-sequenced inbred lines in the University of Kansas greenhouse under standard conditions (Monnahan and Kelly 2015) in three different cohorts. For each cohort, we grew more plants than needed for tissue collection and randomly selected plants for sampling soon after germination. We chose a recognizable and consistent stage at which to collect tissue, which we call the late floral bud stage. These are unopened flower buds (approximately 2-6 mm in length) on the first flowering node (so the corolla is presumably not fully expanded), but are advanced enough that buds on the second flowering node are visible. We chose bud tissue to enrich for transcripts related to flower size. When beginning the first cohort, it was unclear if this tissue type/amount would yield enough RNA for adequate sequencing. We thus pooled bud tissue from three plants of the same line in each tube prior to RNA extraction. Biological replicates were then multiple tubes of pooled tissue, all from the same line. Pooling was done randomly with regard to flowering time (ie. if 6 plants per line were sequenced, 3 in each of 2 tubes, one tube was not all three earliest flowering plants). This process was not repeated in cohorts 2 and 3, for which 1-3 biological replicates (plants) of each line were collected and sequenced separately. We collected tissue into liquid nitrogen at the same time of day (with regard to both actual time and hours after greenhouse lights turn on) within a two-hour window that was consistent between cohorts.

We ground the collected tissue finely with a plastic micropestle and extracted RNA using the Qiagen RNeasy Plant Mini Kit (Hilden, Germany). We generated sequencing libraries using the QuantSeq 3’mRNA-Seq Library Prep Kit for Illumina (Lexogen, Vienna, Austria) per protocol, modified to perform half reactions, and we sequen ced the libraries using NextSeq HO-SR75bp (Illumina, San Diego CA, USA) at the University of Kansas Genome Sequencing Core. Each cohort was sequenced separately with a maximum of 96 samples per flow cell (a total of 4 flow cells and 281 individual samples).

To calculate read counts, we implemented the programs in Lexogen’s BlueBee pipeline. First, we trimmed reads with bbduk (k=13, ktrim=r, useshortkmers=t, mink=5, qtrim=r, trimq=10, minlength=20) from BBTools 38.86 (Bushnell 2014) and aligned reads to the *M. guttatus* V2.0 reference genome (Phytozome, Hellsten et al., 2013) with STAR 2.5.0a (Dobin et al. 2013) using Lexogen’s recommended parameters (outFilterMultimapNmax 20, alignSJoverhangMin 8, alignSJDBoverhangMin 1, outFilterMismatchNmax 999, outFilterMismatchNoverLmax 0.1, alignIntronMin 20, alignIntronMax 1000000, alignMatesGapMax 1000000). Finally, we counted transcript copies using htseq-count 0.11.2 and the genome annotation (Anders et al. 2014). The output is a table of read counts for each transcript. We then removed 5 samples that had fewer than 250k mapped reads (mean for remaining samples of 3,877,524 mapped reads) and normalized the counts for each sample (to account for variable library quality and sequencing depth) using the estimateSizeFactors function in DESeq2 1.28.1 (Love et al. 2014).

### Predicting transcript levels from SNPs

Across all samples, 28,615 of 33,573 total annotated transcripts had at least one mapped read. We kept each gene isoform as a separate transcript. We first filtered out any transcripts which had mapped reads in fewer than 5% of samples, and which did not have 10 or more mapped reads in at least one sample. This left 20,463 transcripts for association mapping. The vast majority of genes (19,721 out of 20,463), had only one isoform with mapped reads. To account for the effect of cohort, we fit a linear model using lm in base R (R Core Team, 2013) to each transcript with cohort as a categorical predictor and then subtracted the estimated effect from each read count. We then transformed each transcript’s expression in each sample by two methods: 1) log(expression + 1), and 2) Box-Cox transformation (Box and Cox, 1964) using the boxcox() function in the R package *EnvStats* 2.3.1 with a range for λ between −5 and 5 (Millard, 2014). Because some counts were negative after factoring out the effect of cohort, we shifted the distributions of all gene counts such that the minimum value was 0 for both types of transformation. Additionally, because Box-Cox transformation cannot accommodate zeros, we added a small value to each count that was equal to 10% of the minimum difference between any two samples (such that the difference between that value and zero was essentially undetectable in the original counts). Finally, we averaged every gene’s expression across plants with each inbred line.

We obtained a filtered set of polymorphisms of the sequenced lines by starting with sites called by Troth et al. (2018). We kept only biallelic SNPs with a minor allele frequency above 2.5% that were called in at least half of the sequenced lines. We then pruned these sites for local LD using PLINK 1.90b3.38 (Purcell et al. 2007) with a window size of 50 SNPs, a step size of 10 SNPs, and an R^2^ threshold of 0.9. This left 2,952,894 SNPs for downstream analysis. We performed the GWAS using gemma 0.98.1 (Zhou and Stephens 2014) by first constructing a centered relatedness matrix using all filtered, but unpruned sites. Using the relatedness matrix, we calculated the ‘chip heritability’ for each gene; the proportion of transcription variation that can be attributed to all SNPs. Finally, we used the univariate linear mixed model (−lmm) in gemma, which in the case of no covariates takes the form: expression = SNP genotype effect + random effect of relatedness + error. We used p-values taken from the likelihood ratio test to find associations between the levels of 20,463 transcripts and each of the 2,953,894 SNPs.

We classified the associations as cis-acting if the site was within 25kb of any part of the transcribed gene and trans-acting otherwise. This distance-based approach for calling cis-effectors can be undermined by long-distance LD. A physically distant SNP (which we would classify as trans) might be associated with expression simply because it is in LD with a cis-acting SNP. For this reason, we excluded data from the meiotic drive locus on chromosome 11 (a known region of extended LD, Fishman and Willis 2005, Fishman and Saunders 2008, Fishman and Kelly 2015) from genome-wide summaries. Overall, the sequenced lines from IM show a rapid decay of LD as inter-SNP distances exceed 10kb (Puzey et al. 2017) which makes our 25kb cutoff conservative. Cis- and trans-acting mutations can be distinguished more directly using allele-specific expression data (Wittkopp et al. 2004, Springer and Stupar 2007, Tirosh et al. 2009, Shi et al. 2012, Osada et al. 2017, Signor and Nuzhdin 2018), but only in heterozygous individuals and here we are measuring expression in highly homozygous inbred lines.

### Predicting phenotypes from gene coexpression modules

To identify sets of coexpressed genes, we used WGCNA 1.69 in R (Langfelder and Horvath 2008) using the sample normalized and cohort factored, but untransformed, counts as input with a power of 3, max block size of 21000, minimum module size of 30, dynamic tree cut method, correlation using dissimilarity, and merge cut height of 0.25. During co-expression analysis, one sample was removed as an outlier. WGCNA identified 37 modules of coexpressed transcripts (Table S1). Next, we extracted the line means for 13 traits from Troth et al. (2018) for the 151 lines used in this study (germination date, days to flower, corolla width, corolla length, floral tube length, throat width, stigma length, anther length, height at flowering, first flowering node, width of widest leaf, and the first two principal components calculated from all floral dimensions. To look for associations between gene expression modules and measured traits, we Box-Cox transformed each module’s eigen expression value (which is the first principal component of a PCA for expression of all member genes in a module), as well as every trait and fit a linear model (all in R, R Core Team, 2013). We used the eigen gene expression for each module as a predictor in a simple regression, as well as fitting the multiple regression for each trait using all 37 modules simultaneously. We also used the program stepAIC from the R package MASS 7.3-52 (Venables and Ripley 2013) to choose a lowest AIC (Akaike Information Criterion) regression model including some but not all modules as predictors.

For each multiple regression model (all modules vs. AIC-chosen set), we used permutation to test for significance. For each trait, we randomly sorted genes into modules with the same number of genes, calculated the eigen gene expression value (PC1) for each module for each line, Box-Cox transformed the module eigenvalues, and then included them in multiple regression. In each case, we fit two models, one with all permuted modules and one with the stepAIC chosen set. We permuted the module composition 1000 times to generate distributions of R^2^ for each trait. Reported p-values are calculated from those distributions. We next used the coefficients from the best-fit model to predict trait variances and covariances. For each trait, we estimated the effect of each module included in the best fit model from a multiple linear regression and constructed an equation to predict trait value for each line:

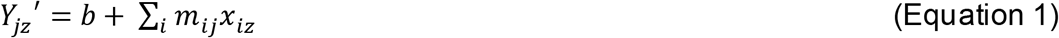

where *Y*_*jz*_′ is the predicted value of trait j for line z, *m*_*ij*_ is the estimated effect of module i on trait j, *x*_*iz*_ is the eigen expression of line z for module i, and the sum is taken over all modules in the model. The covariance of predicted values for traits j and k is:

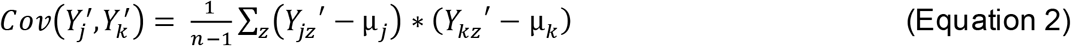

where μ_*j*_ is the mean of trait j, μ_*k*_ is the mean of trait k, and the sum is taken over all n lines. Calculations were done using a custom python script (Supplemental File 1). We permuted traits against module values for testing. For each permuted set, we again found a best-fit model with a subset of gene expression modules and asked how much trait covariation we could predict (using the above method) to generate a distribution. We determined if the amount of trait covariation explained by the real gene expression data, as represented by modules, was significant using alpha levels calculated from the permuted distribution.

Gene expression data has been submitted to NCBI’s SRA (submission number SUB9780070). Python script used to generate trait covariances is available as Supplemental File 1.

## Results

### Genetic effects on transcription levels of individual genes

The number of detected associations between genotype and expression, as well as the putative type of regulatory association (cis vs. trans), are contingent on how we transform transcript counts. Before testing for genetic effects on gene expression, we transformed expression read count data using two methods, log(count + 1) and Box-Cox with λ estimated for each gene separately. Using a rounded p-value cutoff of 1e-12 (Bonferroni = 8.27e-13), we identified 106,585 significant SNP/transcript associations using log(counts+1) transformed counts (10,087 cis associations and 96,498 trans, Fig 1A). Using Box-Cox transformed counts, over 90% of the significant trans-effects evaporate and we find only 8,088 cis and 7,685 trans associations (Fig 1B).

**Figure 1.**
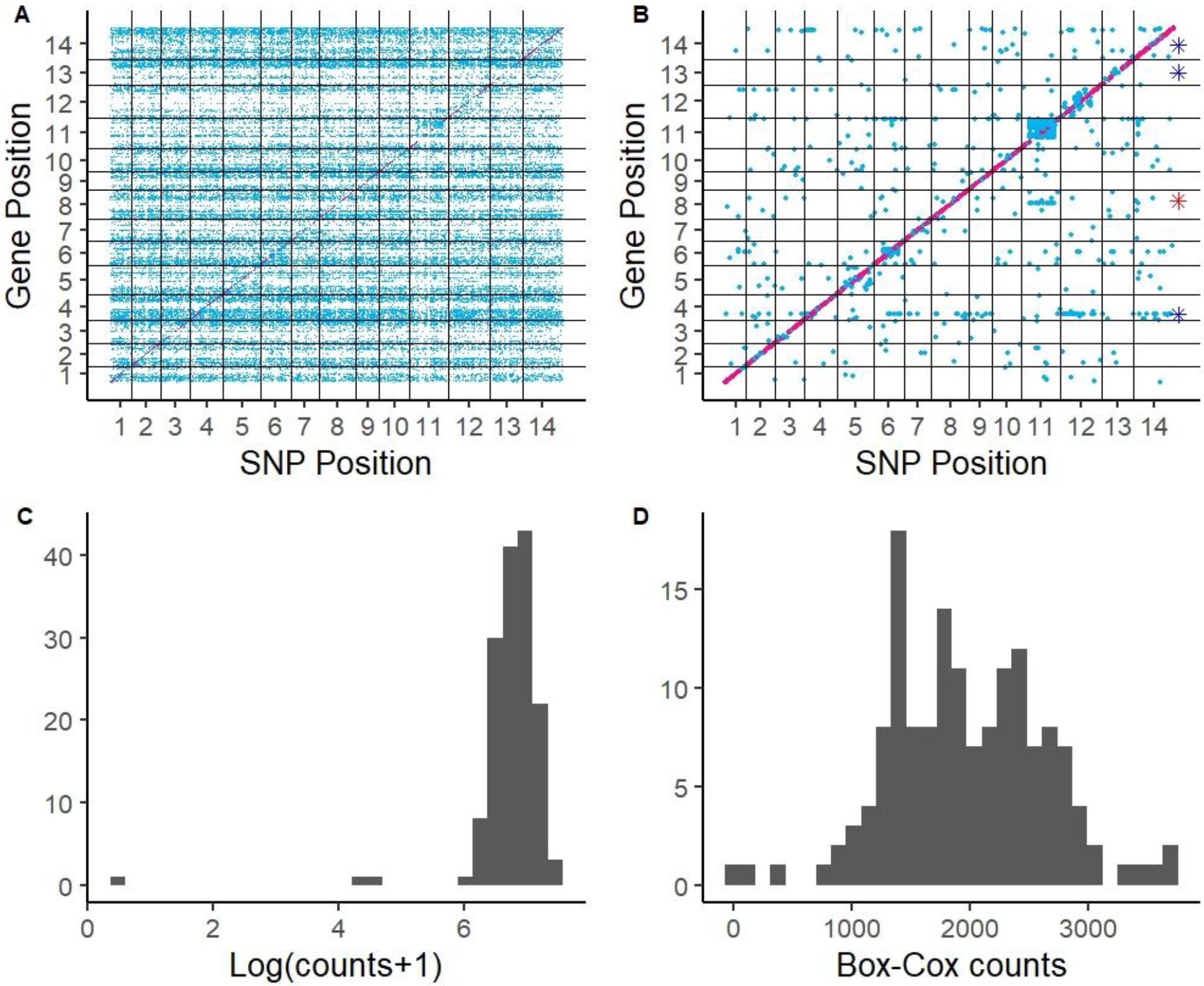
Genome-wide associations of gene expression. Above: Transcript levels were normalized using (A) log(count+1) or (B) Box-Cox. Associations are designated as cis (pink) or trans (blue). Chromosomes are numbered along both axes. Points are larger in panel B to aid visualization. The putative misassembly is indicated with a red asterisk. The three genes with the most transassociations are indicated with blue asterisks. Distributions of read counts for a representative gene, Migut. D00004, which had 3,665 SNP associations with Log(counts+1) transformation (C), but none with Box-Cox transformation (D).

A careful inspection of the differences between the two methods indicates that the Box-Cox results are more reliable. With log(counts+1) transformation, individual genes often have skewed distributions with a small number of lines exhibiting atypically high or low expression (Fig 1C, Fig S1). The few outlier lines with extreme expression will harbor the same minor (and in most cases very rare) allele at many loci. In fact, the majority of the 96,498 trans-regulatory associations (Fig 1A) involve SNPs with a minor allele frequency (MAF) between 2.5-5% (Fig 2A). When unlinked but rare alleles occur together in the same lines, LD is high owing to “rarity disequilibrium” (Houle and Márquez, 2015). If those same lines have extreme expression, all of the linked SNPs will show a strong association with expression.

**Figure 2.**
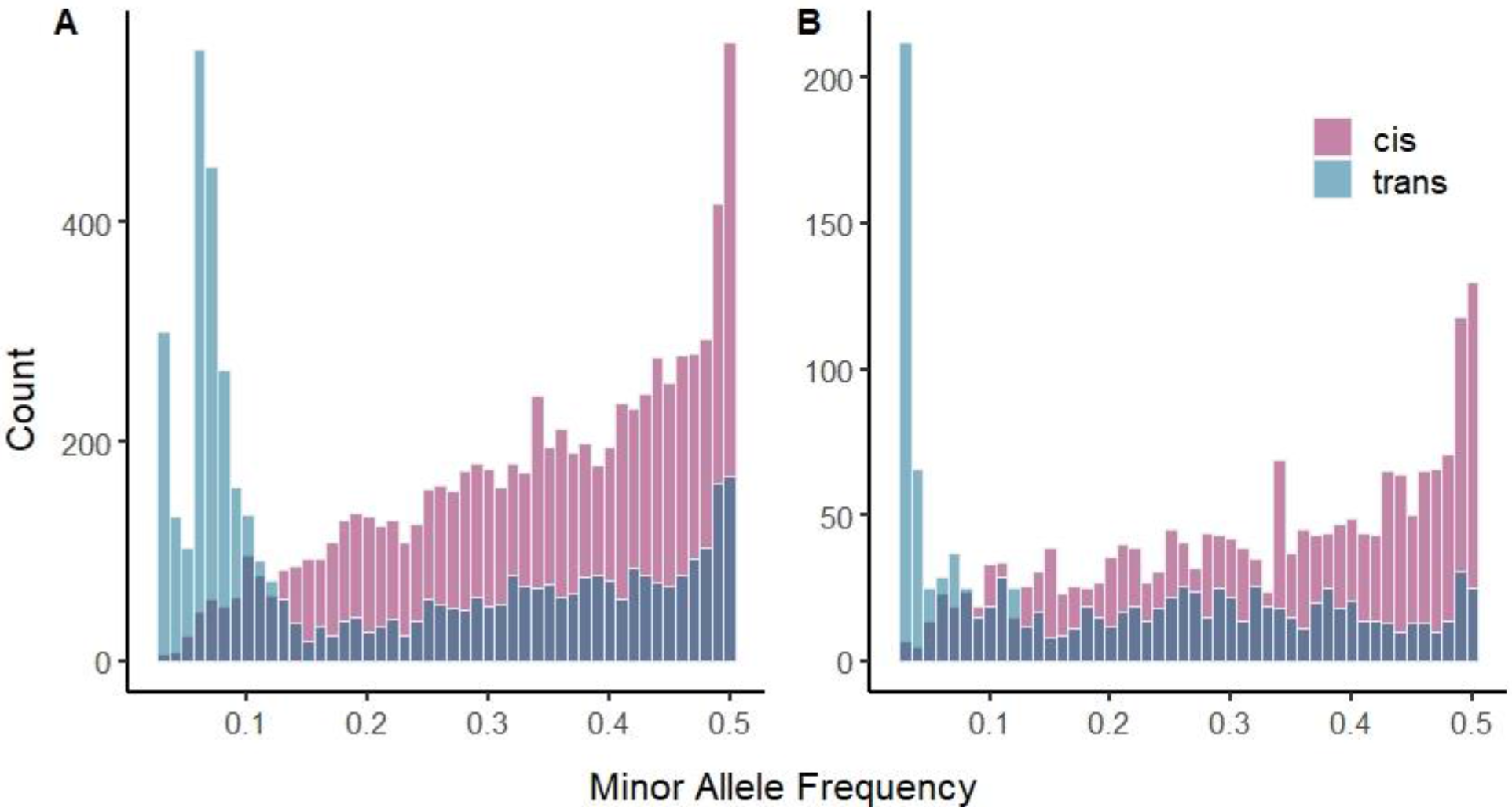
(A) Minor allele frequency distribution for all associations excepting Chromosome 11. (B) The subset of associations in the top quartile of effect sizes for each regulatory category. Cis- acting variants in pink and trans-acting variants in blue.

The Box-Cox transformation provides a scale adjustment specific to each gene. In transcripts with the largest number of genetic associations in the log(counts+1) analysis, Box-Cox more completely “normalizes” expression reducing the effect of outliers (Fig 1C-D, Fig S1). Estimates from the Box-Cox are less affected by the pull of extreme values and we will subsequently limit attention to these tests. The mean estimated heritability of gene expression was 0.31 (see Fig S2 for the full distribution). For genome-wide analyses, we removed SNPs on chromosome 11 because the large block of apparent trans-effects on chromosome 11 are within the meiotic drive locus (Fishman and Willis 2005, Fishman and Saunders 2008, Fishman and Kelly 2015). The Drive allele is essentially a single DNA sequence over >5Mb of DNA segregating in the inbred lines at ~30%. As a consequence, it is impossible to distinguish trans-acting SNPs from those that are simply in linkage disequilibrium with cis-acting SNPs. This leaves 7,832 cis and 4,626 trans associations.

Significant trans-effectors have a larger average effect size (cis=0.54, trans=1.16, F-value = 9.07, p-value =2.6e-3), and effect size is negatively correlated with MAF (both log-transformed; estimated effect of cis-SNP effect size on MAF = −0.029, t = −8.347, p < 2e-16; estimated effect of trans-SNP effect size on MAF = −0.025, t = −4.351, p = 1.39e-05; Fig S3). Since we find trans-acting SNPs have a distribution skewed toward low frequency, it follows that such mutations would also have larger effect sizes, as has been reported in other systems (Josephs et al. 2020). Across the genome, the distribution of minor allele frequency differs greatly between cis- and trans-acting mutations (Fig 2A). We find many rare alleles responsible for trans-regulatory effects on gene expression, and an increasing number of cis effects at higher minor allele frequency (MAF). We find a mean MAF for cis and trans associations of 0.342 and 0.215, respectively (F-value = 2197, p-value < 2.2e-16). The MAF distributions for cis- and trans-acting sites are both different from the MAF distribution of the entire genome (Fig S4, two-sample Kolmogorov-Smirnov tests: cis-to-all comparison: D = 0.51866, p < 2.2e-16, trans-to-all comparison: D = 0.1661, p < 2.2e-16), and are different from each other (D = 0.43651, p < 2.2e-16). The difference in MAF could be due to differential power to detect cis- vs. trans-acting loci with different effect sizes, since we found larger effect sizes for trans effectors. To test whether differences in effect size were driving the MAF pattern, we took only the top quartile of effect sizes (after normalizing by mean expression level) in each regulatory class. The pattern remains the same (Fig 2B) – an excess of common cis-acting alleles and an excess of rare trans-acting alleles. Finally, we established that the pattern is insensitive to our distance cutoff (25kb) for trans-effectors. If we limit trans to SNPs that affect expression on different chromosomes (Fig S5), the cis/trans difference remains.

We find no evidence for “trans-eQTL hotspots”, single SNPs affecting many genes. In fact, there are more genes with transcript levels that are affected by many trans-SNPs than SNPs with more than 2 trans-associations (Fig S6). The three genes with the most trans-acting SNPs are Migut.D00926 (160 SNPs) annotated as a jasmonate ZIM domain-containing protein (JAZ); Migut.M00568 (114 SNPs) annotated as a chlorophyll A/B binding protein; and N01403 (105 SNPs) annotated as an auxin-responsive F-box transport inhibitor response protein. For genes M00568 and N01403, the trans-associations are concentrated on the same chromosome as the gene (Fig 1B). There is an apparent association between the Chr11 Drive Locus and one gene on chromosome 8 (Fig 1A). This gene (Migut.H01175) has three putative homologs in the *Mimulus* genome, only one of which was expressed in our samples (Migut.K01148). We found that all of our samples had high expression for only one of the two genes (Fig S7), and low to no expression for the other, which could indicate mis-assembly. Indeed, when we mapped reads from two samples with expression of either gene to the newer reference genome build (*Mimulus guttatus* TOL v5.0, DOE-JGI, http://phytozome.jgi.doe.gov/) they all mapped to the same region on chromosome 11, which supports that Migut.H01175 is a mis-assembled isoform of Migut.K01148. Finally, the number of cis-associations for a gene is positively correlated with gene size (effect estimate for log(number of associations+1) ~ log(gene length) = 0.1797, t-value = 6.04, p = 1.87e-9 Fig S8).

Rare allele load refers to the proportion of segregating loci at which an individual carries the minor allele, if the population frequency of that allele is very low. It is similar to the concept of deleterious mutation load, but assumes nothing about the fitness effect of individual rare alleles. Instead, it is usually used to test whether or not there is a cumulative fitness effect of harboring many rare variants. This load predicts dysregulation of gene expression in maize (Kremling et al. 2018) and the severity of inbreeding depression in *M. guttatus* (Brown and Kelly 2020). We tested whether lines with an excess of rare alleles exhibit differing patterns of expression, but found no correlation between load and the number of genes showing extreme expression (plus/minus two standard deviations from the mean, Fig S9). We also find no clustering of lines by rare allele load in gene expression principal component (PC) space (Fig S10). The many associations between gene expression and rare variants suggested by Fig 1A (and by the MAF of associations removed by Box-Cox transformation) is thus likely not a real cumulative effect of many rare alleles generating extreme gene expression genome-wide.

### Construction of gene co-expression networks

We next sought to establish sets of genes that co-vary in expression across inbred lines. Using the cohort and individual normalized gene expression counts, WGCNA identified 37 modules of co-expressed transcripts. Each module includes between 37 and 5767 genes (mean 553, median 231) and each transcript (gene) belongs to only one module (Table S1). WGCNA groups genes with correlated expression and further collapses groups such that gene expression between modules should not be highly correlated (R^2^ > 0.8). However, eigengene expression (principal component 1 for the PCA of all genes in a module) of a few pairs of modules remain moderately correlated (20 of 666 pairwise comparisons with 0.74 > R^2 > 0.5) (Fig S11).

To determine if the apparent purifying selection on trans-effecting sites (Fig 2A) is due to their impact on regulatory networks, we calculated the “connectedness” of each gene by correlating the gene’s expression with the eigengene expression value of its module. This measures how predictive a gene’s expression is of the expression of all genes in the module. The distribution of correlation coefficients (R^2^ with module PC1) is highly right-skewed (Fig 3A). For this reason, we grouped genes by “connectedness” quartile and then calculated the average MAF of sites affecting each gene either in cis or in trans. We find a consistent difference in MAF of cis- and trans-effectors, with trans having lower MAF, especially in the highest quartile for “connectedness,” for which MAF is significantly lower than in all other categories (effect estimate for quartile 4 on trans-MAF = −0.143, p = 1.53e-14, effect estimate for quartile 4 on cis-MAF = −0.022, p = 0.00872) (Fig 3B). We did not find enrichment for any GO terms in the set of genes in connectedness quartile 4, using the closest *Arabidopsis thaliana* putative homologs.

**Figure 3.**
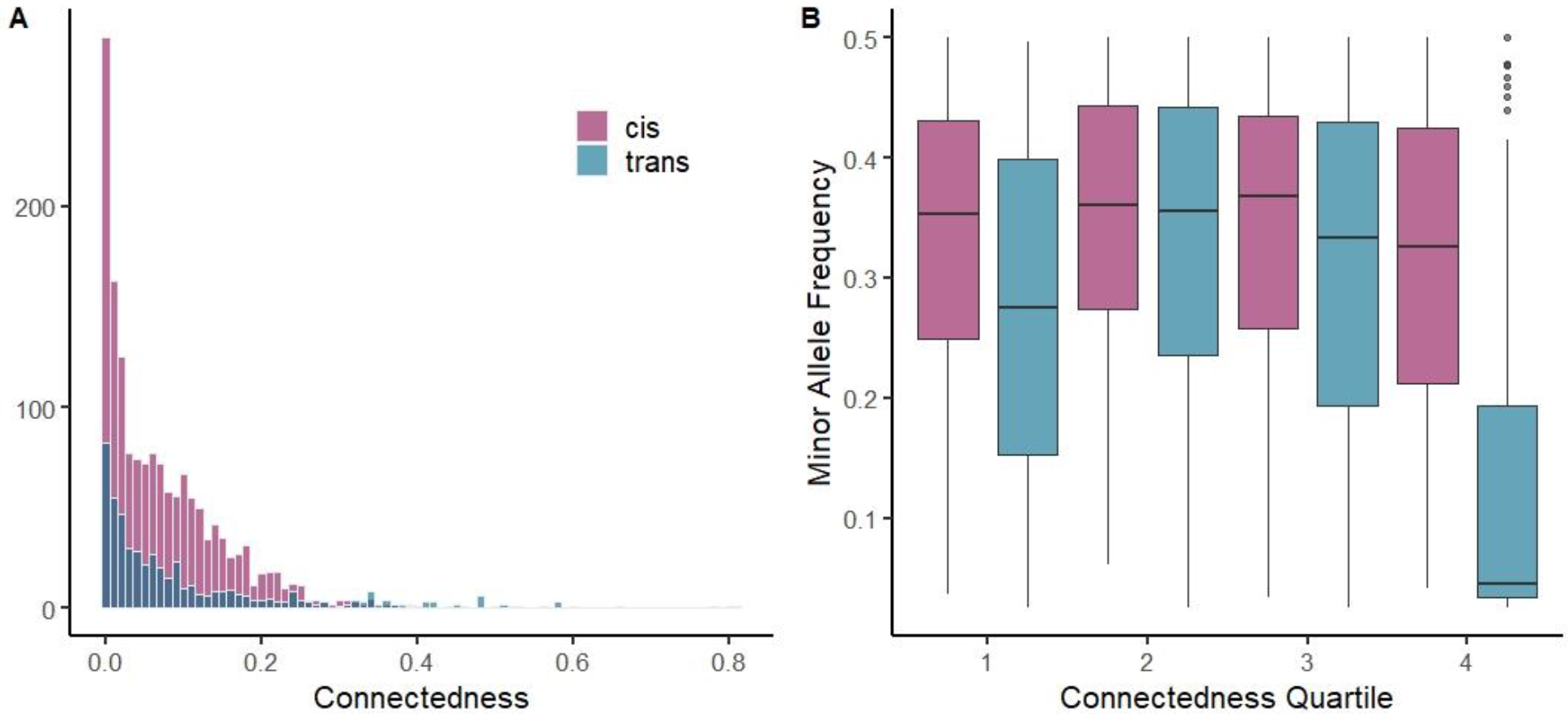
Purifying selection on trans-effectors of highly connected genes. (A) The distribution of connectedness (as measured by R^2^ between a gene and its module expression) for genes with associated cis (pink) and trans (blue) acting variants. (B) The average minor allele frequency of sites affecting each gene in a given connectedness quartile, separated by cis- and trans-acting variants. Each data point in (B) is a gene, which is assigned a quartile and the MAF of sites affecting it is calculated and plotted on the Y-axis.

### Predicting phenotypes from expression

We next tested whether floral bud gene expression affects quantitative traits (Line means from Troth et al. 2018). We use modules of co-expressed genes as predictors of phenotype because this provides a tractable way to incorporate the whole transcriptome. We used multiple linear regression including all 37 modules as predictors of trait, and then chose the AIC-best model for each trait. This selected model included from 6 (widest leaf) to 20 (flower size PC1) of the 37 total expression modules (Table 1). The best-fitting model for each trait explained from 23% to 47% of trait variation, with the strongest prediction being for overall flower size (PC1 in Table 1). To establish statistical significance for prediction of trait variation, we permuted the data in two ways:

**Table 1.**
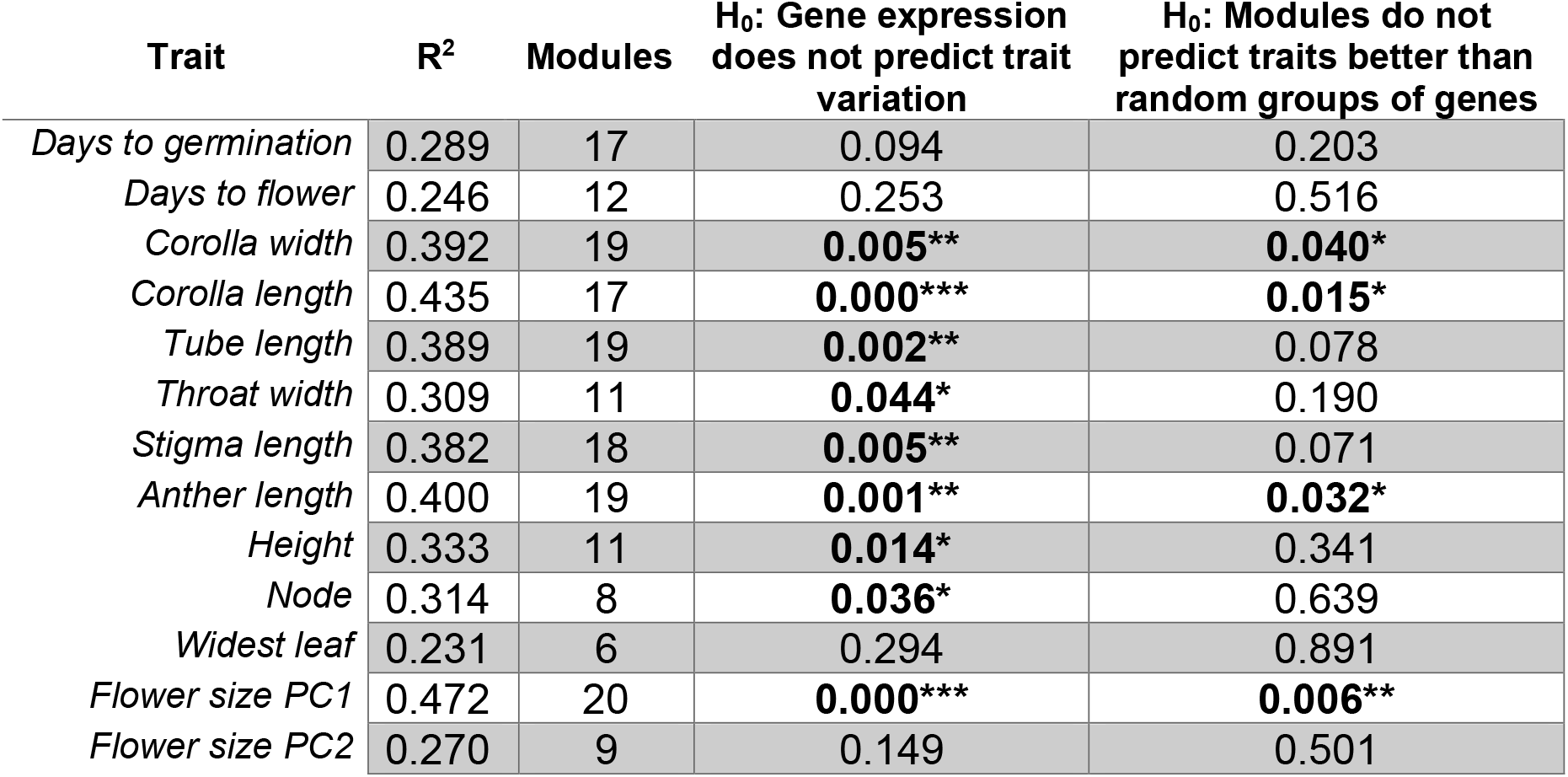
Variation in traits explained by the AIC best-fit model of gene expression modules. Significance was established by permuting either the module eigen values across lines, or the module gene composition. Bold indicates significance : *: p < 0.05, **: p < 0.01, ***: p<0.001.

| Trait | R<sup>2</sup> | Modules | H<sub>0</sub>: Gene expression does not predict trait variation | H<sub>0</sub>: Modules do not predict traits better than random groups of genes |
| --- | --- | --- | --- | --- |
| *Days to germination* | 0.289 | 17 | 0.094 | 0.203 |
| *Days to flower* | 0.246 | 12 | 0.253 | 0.516 |
| *Corolla width* | 0.392 | 19 | **0.005\*\*** | **0.040\*** |
| *Corolla length* | 0.435 | 17 | **0.000\*\*\*** | **0.015\*** |
| *Tube length* | 0.389 | 19 | **0.002\*\*** | 0.078 |
| *Throat width* | 0.309 | 11 | **0.044\*** | 0.190 |
| *Stigma length* | 0.382 | 18 | **0.005\*\*** | 0.071 |
| *Anther length* | 0.400 | 19 | **0.001\*\*** | **0.032\*** |
| *Height* | 0.333 | 11 | **0.014\*** | 0.341 |
| *Node* | 0.314 | 8 | **0.036\*** | 0.639 |
| *Widest leaf* | 0.231 | 6 | 0.294 | 0.891 |
| *Flower size PC1* | 0.472 | 20 | **0.000\*\*\*** | **0.006\*\*** |
| *Flower size PC2* | 0.270 | 9 | 0.149 | 0.501 |

#### Permutation 1

Does gene expression predict trait variation? To test the hypothesis that a model using gene expression explains no more trait variation than by chance, we permuted modules by line. Correlations among traits and among modules were preserved, but randomly associated with each other across lines. Using this method, trait variation predicted by the real gene expression modules is highly significant (p < 0.01) for 7 of 8 flower-size measurements (except flower size PC2), and marginally significant (0.01 < p < 0.05) for height and node (Table 1). These 9 traits are significantly correlated with each other, except for throat width with node (Fig 4, Table S2).

**Figure 4.**
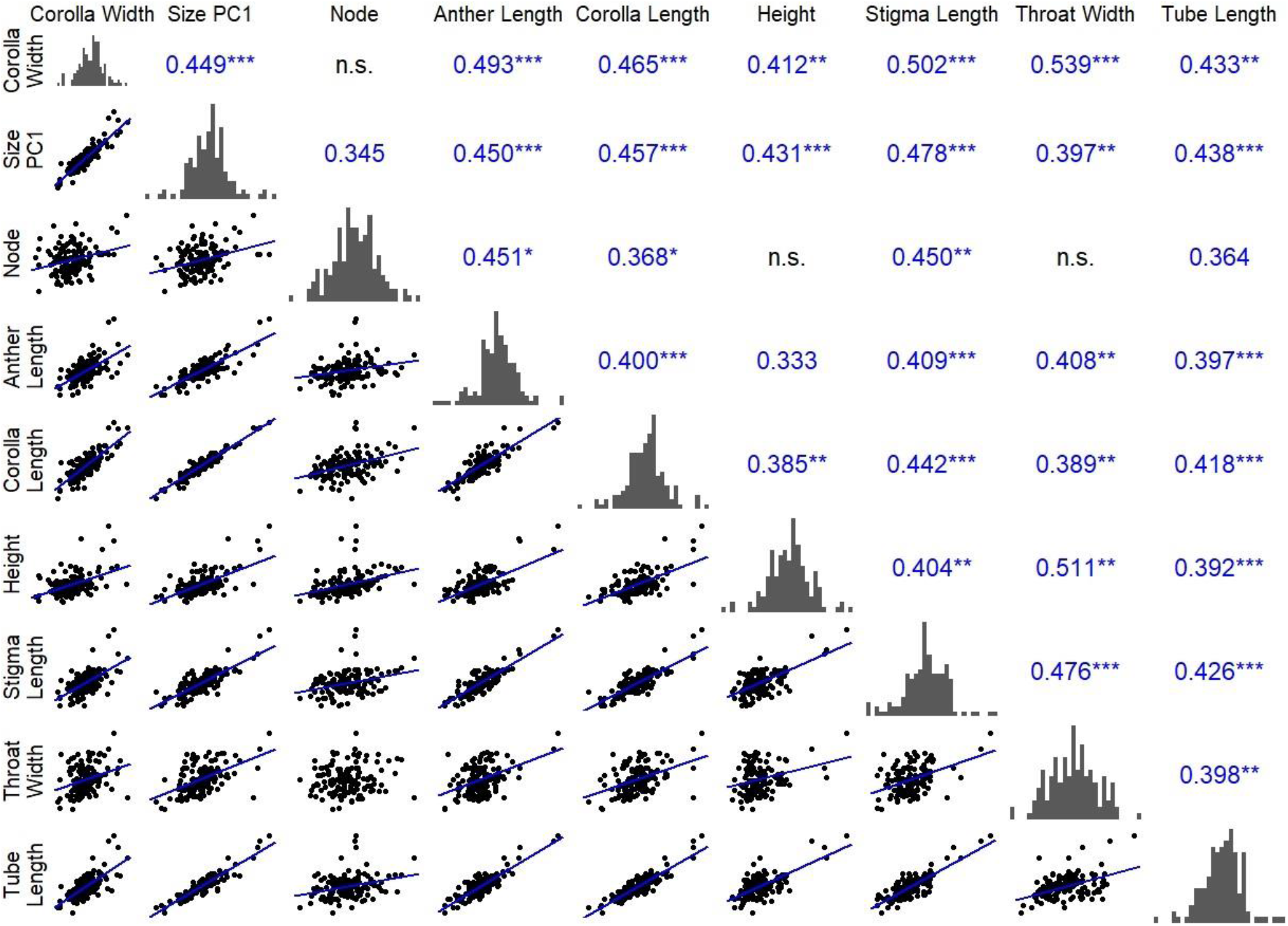
Gene expression predicts trait covariances. The bottom triangle shows trait correlations. A line denotes a significant correlation in a linear regression at p < 0.05. The diagonal displays normalized trait histograms. Fraction covariance explained by gene expression, using the best-fit model coefficients for prediction, is shown in the top diagonal. Any displayed number is significant, asterisks denote levels of significance (determined by permutation, p < 0.05, p* < 0.025, p** < 0.01, p*** < 0.001).

#### Permutation 2

Do gene co-expression modules better predict traits than random groups of genes? The significant prediction of traits by modules does not imply that modules are necessarily the best summary of gene expression for trait prediction. In order to test the hypothesis that the predicted trait variation is just a function of including the whole transcriptome (by creating groups of genes as predictors), we permuted module membership by shuffling genes into random groups of the same size as the real modules. These groups contain the same amount of information in terms of fraction of transcriptome included, but eliminate clustering of genes based on co-expression that defines the real modules. By permuting gene module membership, we find that only 4 traits (corolla width and length, anther length, and flower size PC1) are significantly better predicted by co-expression modules than by random assortment of genes (Table 1). For all other traits, the amount of trait variation explained is attributable to the inclusion of the whole transcriptome, not variation in co-expressed groups of genes. However, while most traits are not significantly better predicted by real modules than scrambled sets of genes, real modules better predict traits than the *average* permuted data set for all but four traits (days to flower, node, widest leaf, and flower size PC2).

Overlapping sets of gene expression modules are included in the best-fit model for the four traits where expression modules are significant by both permutation tests (corolla width and length, anther length, and flower size PC1). All four are significantly predicted (in their own best-fit regression models) by 14 common modules. These traits are all positively correlated (Fig 4). The 14 modules are not correlated (Fig S11), but they affect all 4 traits in the same direction. As a consequence, trait correlations can emerge from the joint effects of uncorrelated modules.

Prior studies indicate that tradeoffs between fitness components (and associated traits) are central to the maintenance of variation in this population (Mojica and Kelly 2010, Scoville et al. 2011, Mojica et al. 2012, Monnahan and Kelly 2015, Monnahan and Kelly 2017, Brown and Kelly 2018). For this reason, we estimated the extent to which gene expression modules generate trait covariances. Among pairwise comparisons between the 9 traits that are significantly predicted by gene expression (Table 1 column 3, and see Permutation 1 above), 35 of 36 pairs are significantly correlated (R^2^ between 0.07-0.91, p < 0.05 for all but node by throat width). We used estimates for the effect of each gene expression module on each trait from the best-fit multiple linear model (Table 1, column 1) to predict trait covariances using equations 1–2. If a module affects two traits, some fraction of the covariance between the traits can be attributed to the shared effect of that module. We find that 26-54% of the covariance between traits is attributable to this module-predicted covariance (35 pairwise comparisons). 33 of 36 covariances are significantly predicted by gene expression modules (Permutation p < 0.05, Fig 4 upper triangle). Modules are most strongly predictive of trait covariances among the four traits that are better predicted by modules than by the randomly grouped whole transcriptome (corolla width, corolla length, stigma length, anther length, and flower size PC1, see Table 1 column 4).

A large fraction of module variation is genetic. For each individual, we calculated the eigen expression (PC1) for each module and tested for an effect of inbred line using an ANOVA. Line explains 57-91% of variance in gene module expression (Table S3). Of the 37 modules, 29 are significantly affected by line (p < 0.05). There is no correlation between the estimated genetic control of a module and the number of traits for which a module is a significant predictor. However, all modules that significantly predict at least half of our measured traits (save one, “brown”) are significantly affected by genotype (p from 0.046 to 8.18e-15, F from 1.34 to 4.04). That is to say, modules that significantly predict many traits exhibit genetic variation among lines.

## Discussion

### Natural selection on regulatory variants

Using a collection of sequenced inbred lines derived from a single natural population of yellow monkeyflower (*Mimulus guttatus*), we have dissected the genetic variation in the floral bud transcriptome. We found 12,458 SNPs with genome-wide significant associations with expression, 62% of which act in cis. Striking differences in the allele frequency spectrum suggest differing selection regimes on cis- and trans-acting regulatory SNPs. Sites proximal to the affected gene are enriched for intermediate frequency variants. SNPs distant from target genes are enriched for rare variants. Hodgins-Davis et al. (2015) argue that gene expression should evolve according to a “house of cards” model, characterized by few mutations with large effects and moderate stabilizing selection (as opposed to a Gaussian model of evolution with many mutations of small effect and weak selection). Stabilizing selection on a quantitative trait with a fixed optimum predicts that minor alleles should be less common than under neutral evolution. Trans-acting mutations are more likely to be deleterious than cis-acting mutations if they have more pronounced effects (see introduction). The results of this study suggest that different selective pressures operate on cis and trans variation, consistent with previous work in a natural population of *Capsella grandiflora* (Josephs et al. 2020). The distribution of MAF for trans-effecting SNPs in Capsella was similar to the Mimulus estimate (Fig 2). However, Capsella exhibits a nearly uniform distribution of MAF for cis-SNPs, while there is a definite inflation of intermediate frequency SNPs in Mimulus. The skew of cis-acting SNPs towards intermediate frequency, relative not only to trans-acting but also the genome as a whole, suggests balancing selection.

We find support for the hypothesis that trans-effectors are routinely subject to purifying selection (Figs 2,3), at least for mutations with large enough effects to be detected in this study. Loci influencing expression in trans can affect multiple components of finely-tuned networks simultaneously. Here, we show that the minor allele frequency of SNPs affecting a gene’s expression is correlated with how well that gene predicts the expression of many other genes (those in the same coexpression module), what we call “connectedness.” Genes that are well-connected in this sense are likely to be the hub of a regulatory network, a role commonly filled by transcription factors (Babu et al. 2004), although we do not detect an enrichment for any particular type of gene in this set. We find that SNPs affecting well-connected genes tend to be lower in frequency and that the magnitude of decrease in MAF is stronger for trans-acting SNPs than cis-acting SNPs (Fig 3). This difference supports the idea that trans-effectors with broad pleiotropic effects on many genes are more likely to affect regulatory hubs and therefore be routinely subjected to purifying selection.

Connectedness of genes affected by trans-SNPs might explain the pattern of purifying selection, but it does not explain why cis-acting variants exhibit an MAF distribution suggestive of balancing selection. Cis-acting variants did have smaller effect sizes, which would explain a difference in severity of purifying selection, but not that allele frequencies at cis- SNPs are more intermediate than the genome-wide average. One potential explanation is that cis-acting variants may evolve on a gene-by-gene basis to counter the pleiotropic effects that trans-acting loci have on many genes. The hypothesis that cis-acting variants might evolve to mitigate trans-pleiotropy is supported by many studies finding opposing cis- and trans effects on the same gene (Coolon et al. 2014, Wang et al. 2015, Mack et al. 2016, Metzger et al. 2017). In this study, we find no such preponderance of compensatory cis/trans pairs. Using a conservative set of 24 genes with both cis-SNPs and inter-chromosome trans-SNPs, we find only one example of cis/trans compensation.

### Genetic effect on traits mediated through gene expression

The MAF distribution (Fig 2) and the relationship between allele frequency and gene connectedness (Fig 3) provide clear evidence for purifying selection on trans-acting regulatory mutations. Cis mutations exhibit a MAF distribution suggesting balancing selection, but this pattern is not clearly explained by cis/trans compensation. A more complete understanding of selection requires that we look at how genetic effects on gene expression translate to effects on phenotypes and, ultimately, on fitness within the natural population. For this, we look to the 13 fitness-related traits measured by Troth et al. (2018; Table 1). Previous field studies, including Troth et al., have shown that these traits are under strong but fluctuating selection in the field (Mojica et al. 2012, Monnahan et al. 2021). We here demonstrate that many of these traits can be significantly predicted by the transcriptome, with the latter abstracted into coexpression modules (Table 1). The most precise prediction is for overall flower size (flower size PC1) and for the component measurements that jointly determine flower size (corolla width and length, stigma and anther lengths).

Stronger prediction of floral traits than for phenology or leaf width is not surprising given that we collected RNA from flower bud tissue. However, the strength of prediction (nearly 50% of variation explained for flower size) is notable given the possibility that flower traits are established far earlier in development (Krizek and Anderson, 2013). This might suggest that trait variation is continually programmed and reinforced through development or simply that gene expression levels might correlate across development time. Prediction precision was likely reduced by the fact that modules were estimated from RNAseq performed on one set of plants, while the mean phenotypes were estimated from different plants of the same inbred lines. Plants from the two experiments almost certainly experienced subtle environmental differences (different greenhouses, growth at different times of the year, different years). The high R^2^ for flower size despite these limitations suggests that stable relationships between genotypes and traits are mediated through transcriptome variation. Additionally, the separation of experiments does avoid a subtle but potentially important bias. When phenotypes and gene expression levels are measured on the same plants, the two can become associated owing to confounding factors, even when there is no effect of expression on phenotype. Imagine that plants differ randomly in receipt of a resource such as soil nitrogen. If nitrogen affects both gene expression and phenotype, expression and phenotype will be correlated even if there is no inherent relationship. Establishing the mean phenotype of each line prior to measuring expression eliminates this source of bias (Rausher 1992).

Gene expression (as collapsed into modules) predicts trait variation and also the covariances between traits (Fig 4). In fact, module predictions account for up to 54% of covariance between traits (throat width and height). Variation in gene expression explains covariances between traits whose correlation is responsible for the life-history tradeoff in fitness components. It has been shown that such a tradeoff can be a mechanism for the maintenance of genetic variation through balancing selection (Brown and Kelly, 2018). As such, the connection to phenotype might be a plausible explanation for the pattern of excess intermediate frequency alleles affecting gene expression, although more work would be necessary to better understand why these loci act overwhelmingly in cis. We also show a substantial genetic component of many of the gene coexpression modules, which indicates that the genetic control of fitness-related traits could be mediated (or potentially compounded) through effects on gene expression.

### Scale of measurement for gene expression

Fig 1 contrasts two different ways to normalize read counts, Box-Cox and log(count+1). The latter is most similar to models typically applied in RNAseq studies, such as generalized linear models that use the log-link function (e.g. DESeq2; Love et al. 2014). When expression is normalized in the same way across all genes (such as with the log(count+1) method), rare alleles occurring in lines with extreme expression produce many false positives as a result of “rarity disequilibrium” (Houle and Márquez, 2015). When counts are instead power transformed using an exponent (λ) estimated for each gene separately (Box-Cox), samples with extreme expression are pulled closer to the mean of the resulting distribution (compare Figs 1C,D). This decreases the occurrence of spurious associations due to rare alleles. We retained the log(count+1) analysis in Fig 1 as a caution for future studies. This issue is likely to emerge in any situation where the absolute count of individuals carrying the rare allele is small (say less than 5).

## Conclusion

The two major findings from this study are connected through our summarization of the transcriptome in terms of gene expression modules. The first result is that cis-acting SNPs tend to have intermediate allele frequencies (relative to the genome as a whole), while trans-SNPs exhibit a rare-alleles model consistent with purifying selection. Trans-acting mutations are most rare if they have broad effects, with the latter measured by how strongly a trans-affected gene predicts the overall expression of its module. The second result is that expression modules predict flower size with a surprising degree of precision. As a consequence, we can attribute substantial fractions of the variance in flower size measures to variation in expression modules (R^2^ values in Table 1). Despite that expression levels of different modules are largely uncorrelated (across lines), they can generate covariances among traits because individual modules influence multiple traits. This ‘transcriptome-explained’ covariance can be a substantial portion of the total covariance across lines (up to 54%, Fig 4).

Our results do not provide a clear explanation for why cis-acting SNPs exhibit allele frequencies consistent with balancing selection, but the prediction of trait covariances suggests future studies that may address this question. Many years of field study of the IM population suggests that trade-offs between traits and fitness components are essential to the maintenance of genetic variation (Mojica and Kelly 2010, Scoville et al. 2011, Mojica et al. 2012, Monnahan and Kelly 2015, Monnahan and Kelly 2017, Brown and Kelly 2018). Plants that develop rapidly have high viability (survivorship-to-flowering) but often produce smaller flowers and can suffer lower fecundity (Troth et al 2018). Fitness trade-offs manifest as genetic correlations between traits like flower size and days to flower. We cannot predict this trait relationship with the current data because the flower-bud transcriptome does not effectively predict days to flower (Table 1). However, the basic modeling approach applied here to intra-floral traits may succeed for development rate if we can sample the relevant tissue at the appropriate developmental stage (or across stages). Future studies that determine the nature and extent of transcriptional control of development rate could provide a more mechanistic understanding of balancing selection.

Gene expression data has been submitted to NCBI’s SRA (submission number SUB9780070). Python script used to generate trait covariances is available as Supplemental File 1.

## Acknowledgements

We thank S. Macdonald, P. Veltsos, L. Hileman, E. Everman, R. Unckless, C. Wessinger, and C. Fiscus for comment on the manuscript, and the Koenig and Seymour labs at UCR for comments on early stages of analysis. JKK acknowledges support from NSF grants DEB-1753630 and MCB-1940785. KEB was supported by NSF PRFB IOS-1907061 and a KU Genome Sequencing Core Award Voucher. This work was supported by the Center for Research Computing at the University of Kansas and the KU Genome Sequencing Core.

## Supplementals

**Figure S1.**
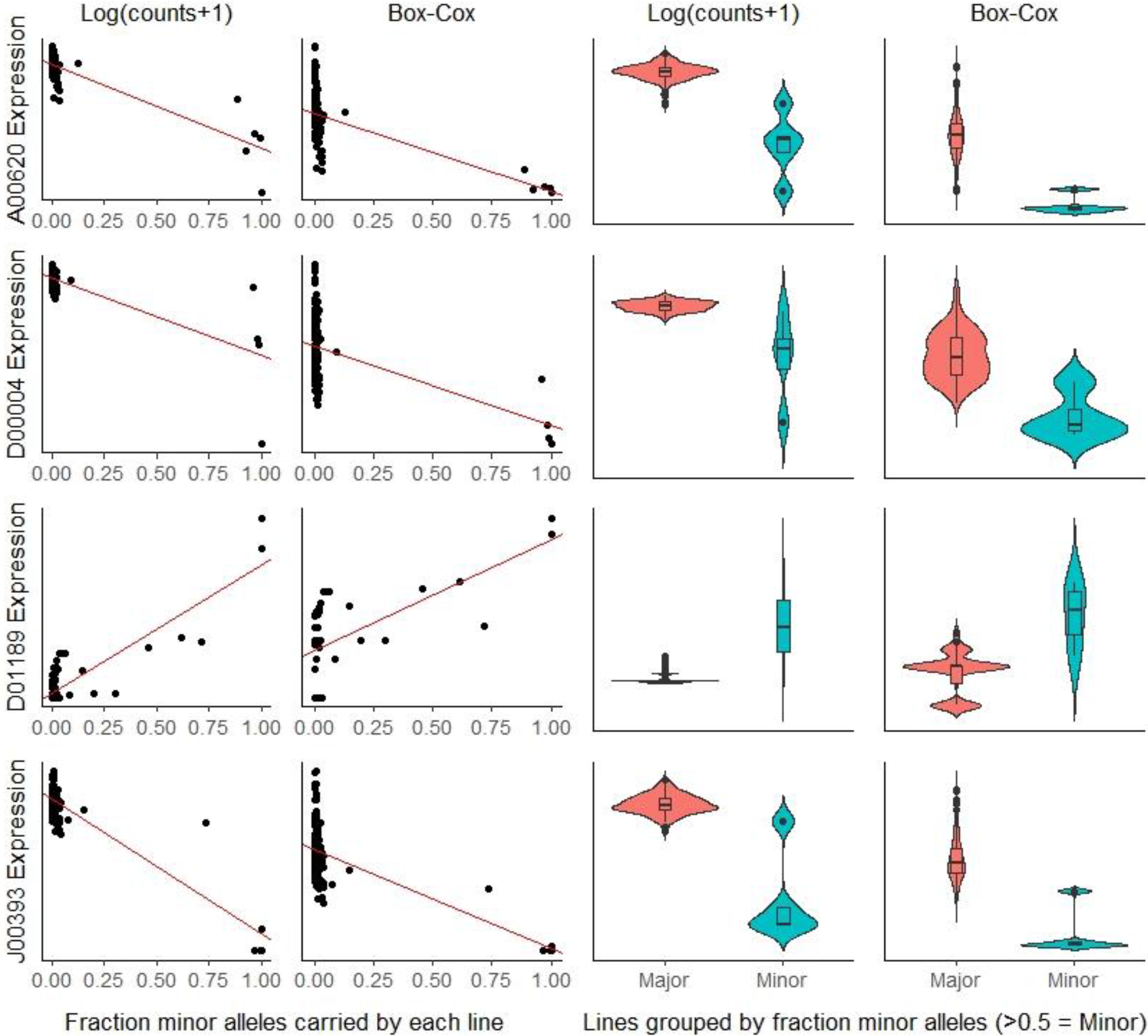
Genes representing how Box-Cox transformation reduces associations caused by “rarity linkage.” The four genes pictured have the most SNP associations of all genes after Log(counts+1) transformation, and no associations after Box-Cox transformation. The first two columns show the relationship between how many *associated* minor alleles a line carries and expression transformed both ways. The second column splits lines into those that carry mostly major vs. minor alleles at associated loci. Note that Box-Cox transformation spreads the distribution among lines carrying the major alleles so that linked rare alleles no longer cause spurious associations.

**Figure S2.**
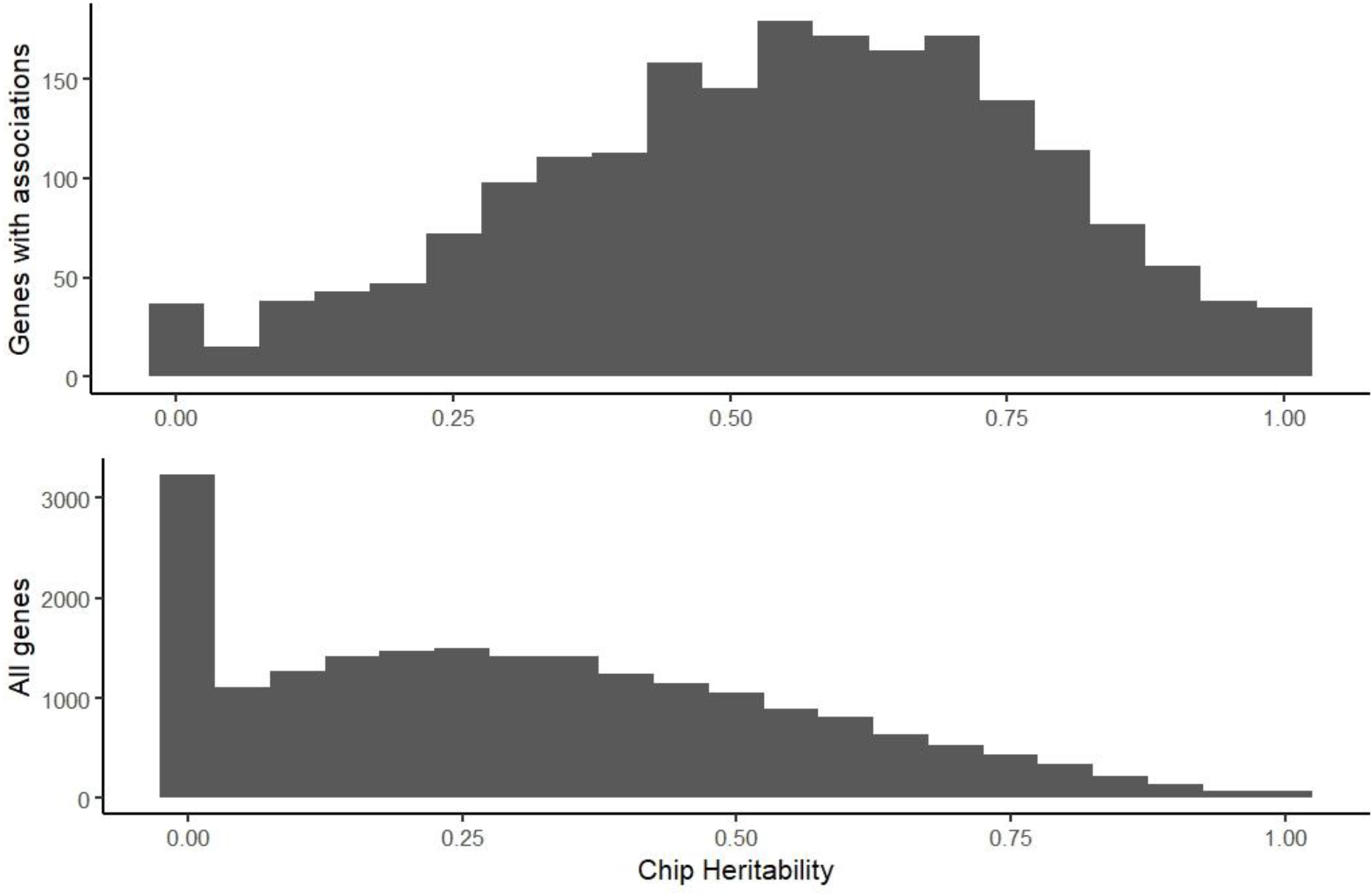
Distribution of chip heritability (proportion of variance in expression explained by all tested SNPs) for all genes tested in the eGWAS (bottom) vs. genes that have significant SNP associations (top).

**Figure S3.**
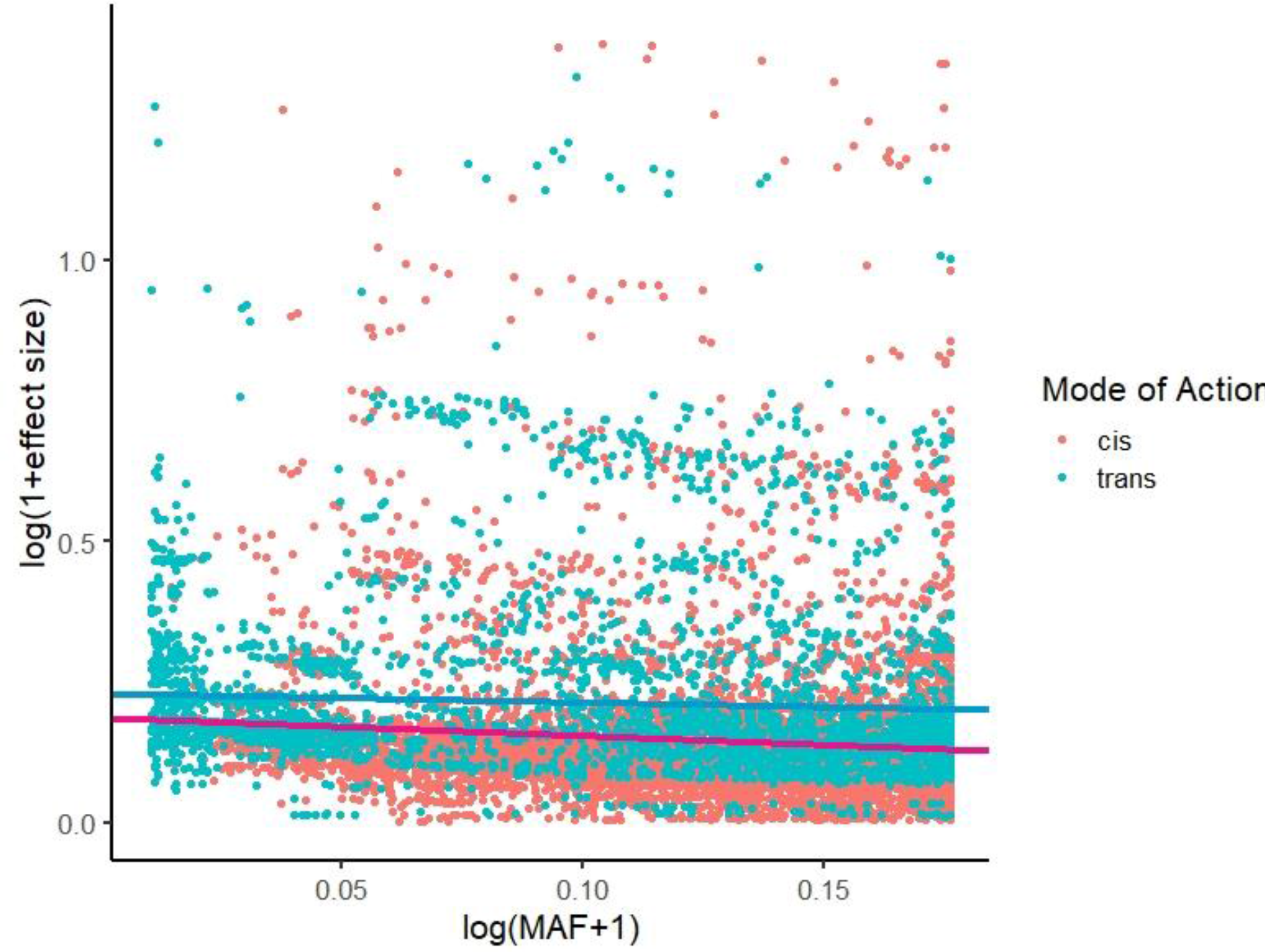
Minor allele frequency of significant SNPs in the eGWAS vs. effect size normalized by mean gene expression. Five extreme outliers were removed. Lines are the estimated correlations, separated by cis vs. trans-acting SNPs (pink for cis, blue for trans).

**Figure S4.**
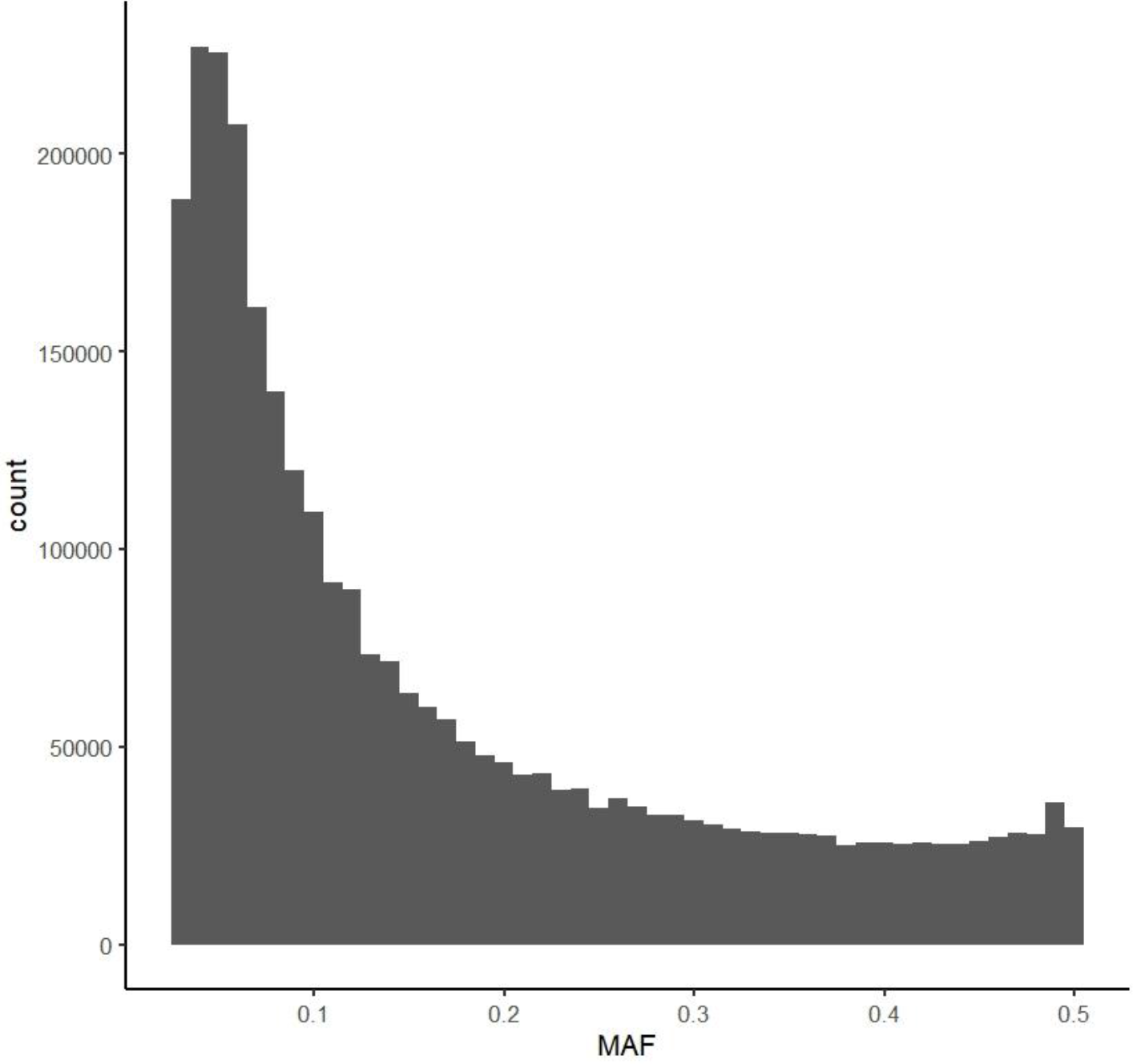
Minor allele frequency (MAF) distribution for all SNPs that were included in testing.

**Figure S5.**
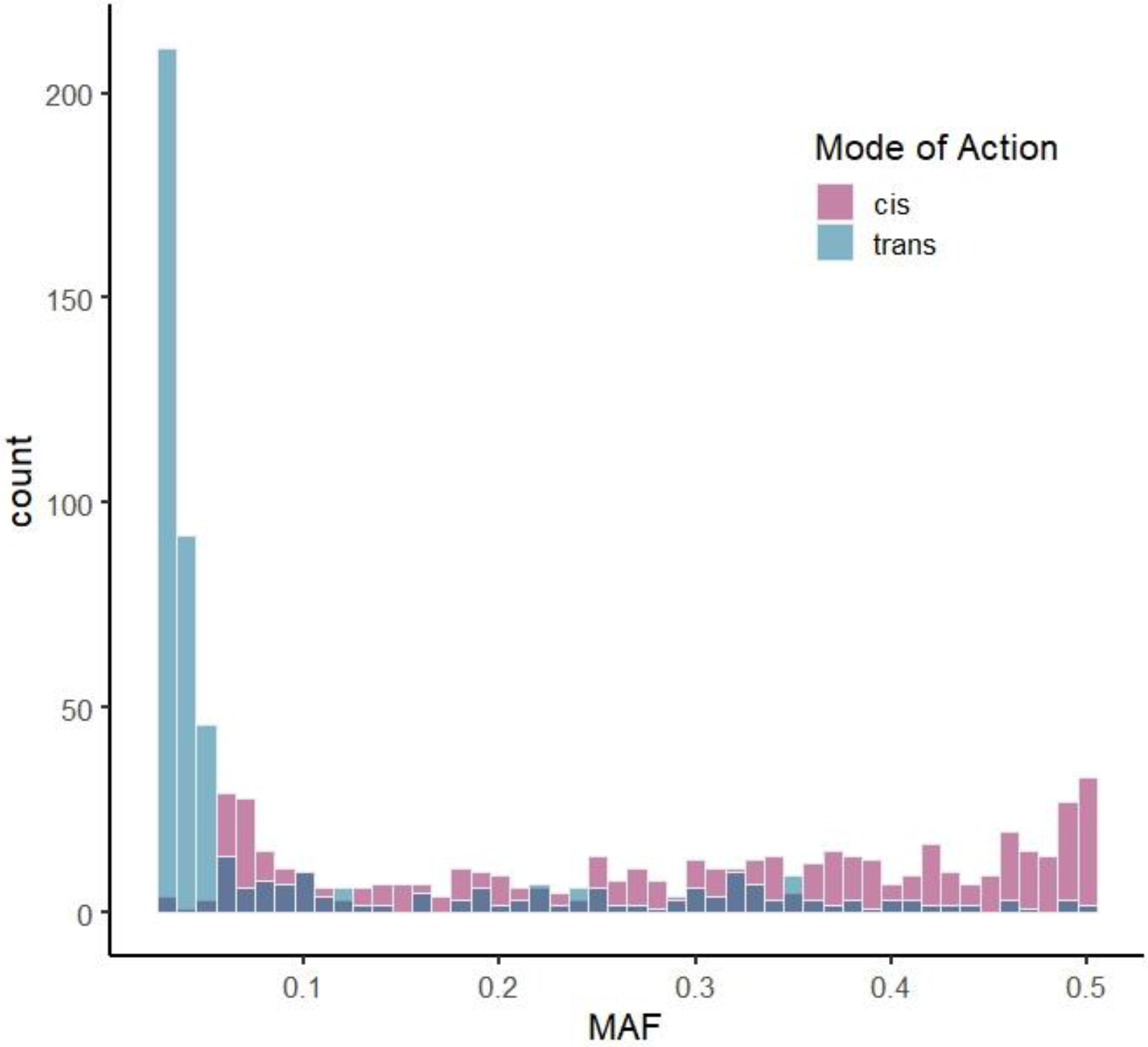
Minor allele frequency distribution of cis- and trans-acting variants when the distance cutoff is relaxed. For these distributions, only SNPs on a different chromosome from the affected gene were classified as trans. Because this generated a substantially smaller set of trans-acting SNPs, the cis-acting SNP set was randomly downsampled to the same number.

**Figure S6.**
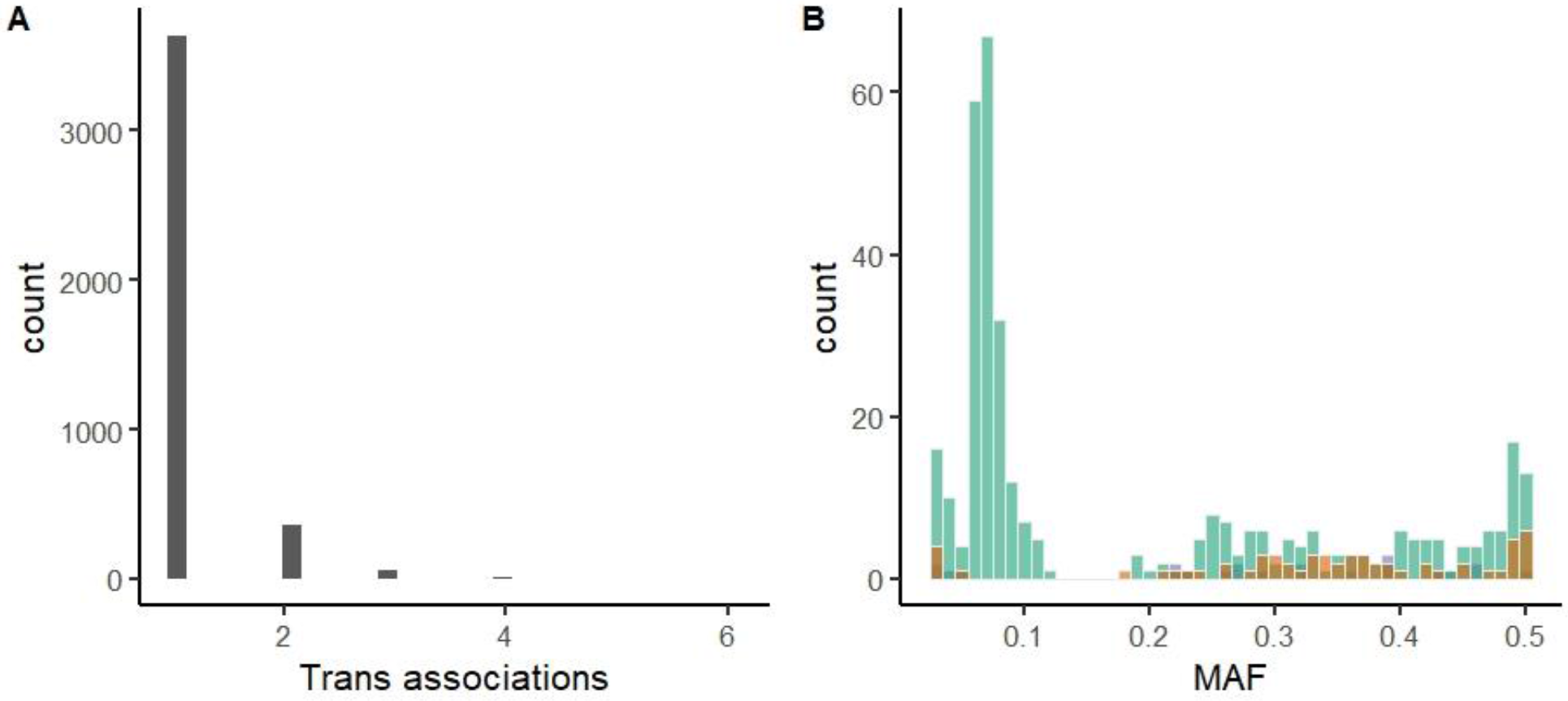
Few SNPs show trans-associations with more than one gene. A) The distribution of the number of genes each SNP affects in trans. B) MAF distribution for sites that affect 2 (bluish green), 3 (orange), or >3 (purple) genes in trans. Note that the distribution is not different from that of all sites with trans-associations.

**Figure S7.**
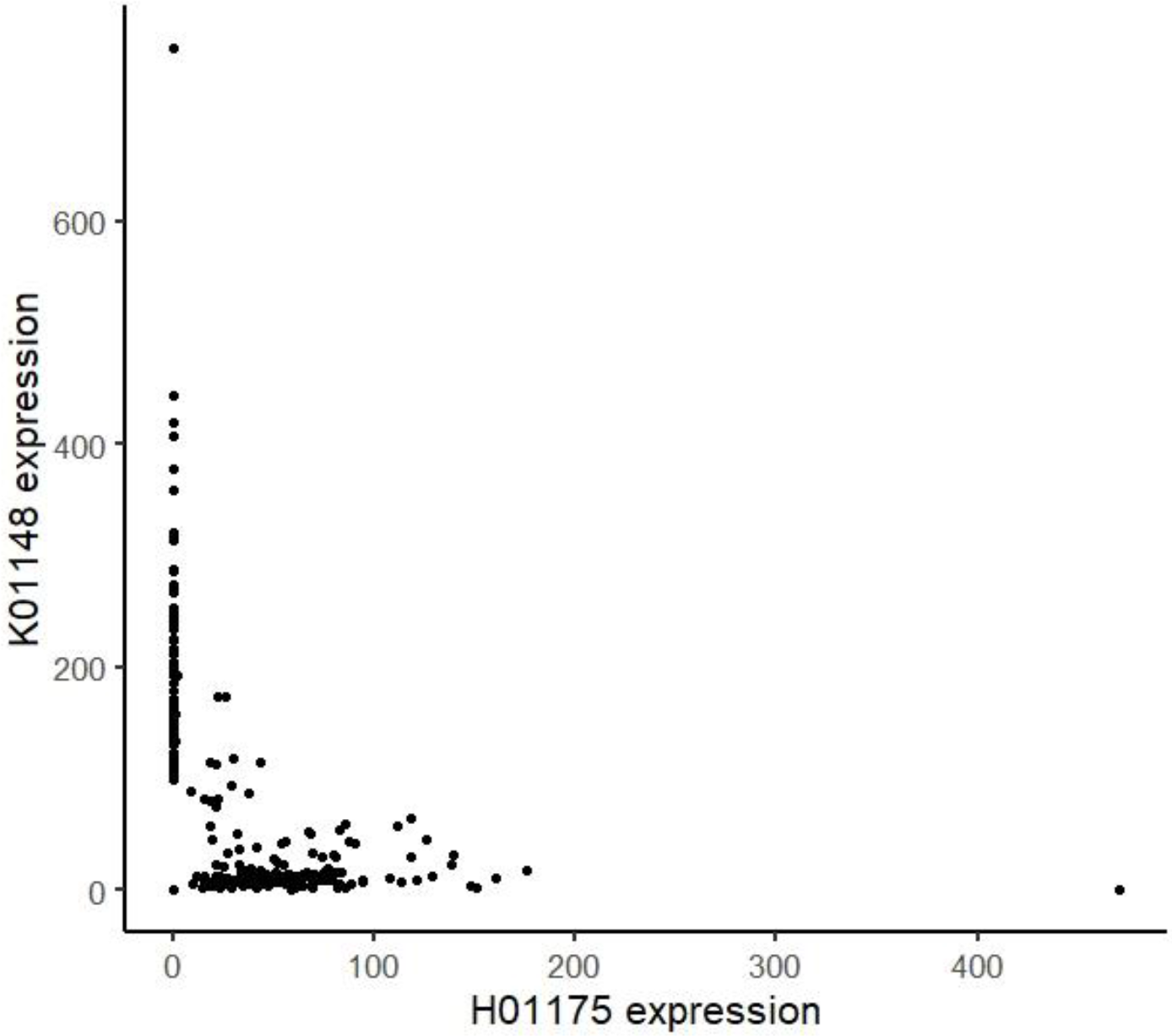
Misassembly of Migut.H01175. For each sample, the apparent expression of Migut.H01175 is plotted vs. expression of Migut.K01148. Reads map to one or the other, indicating misassembly of isoforms.

**Figure S8.**
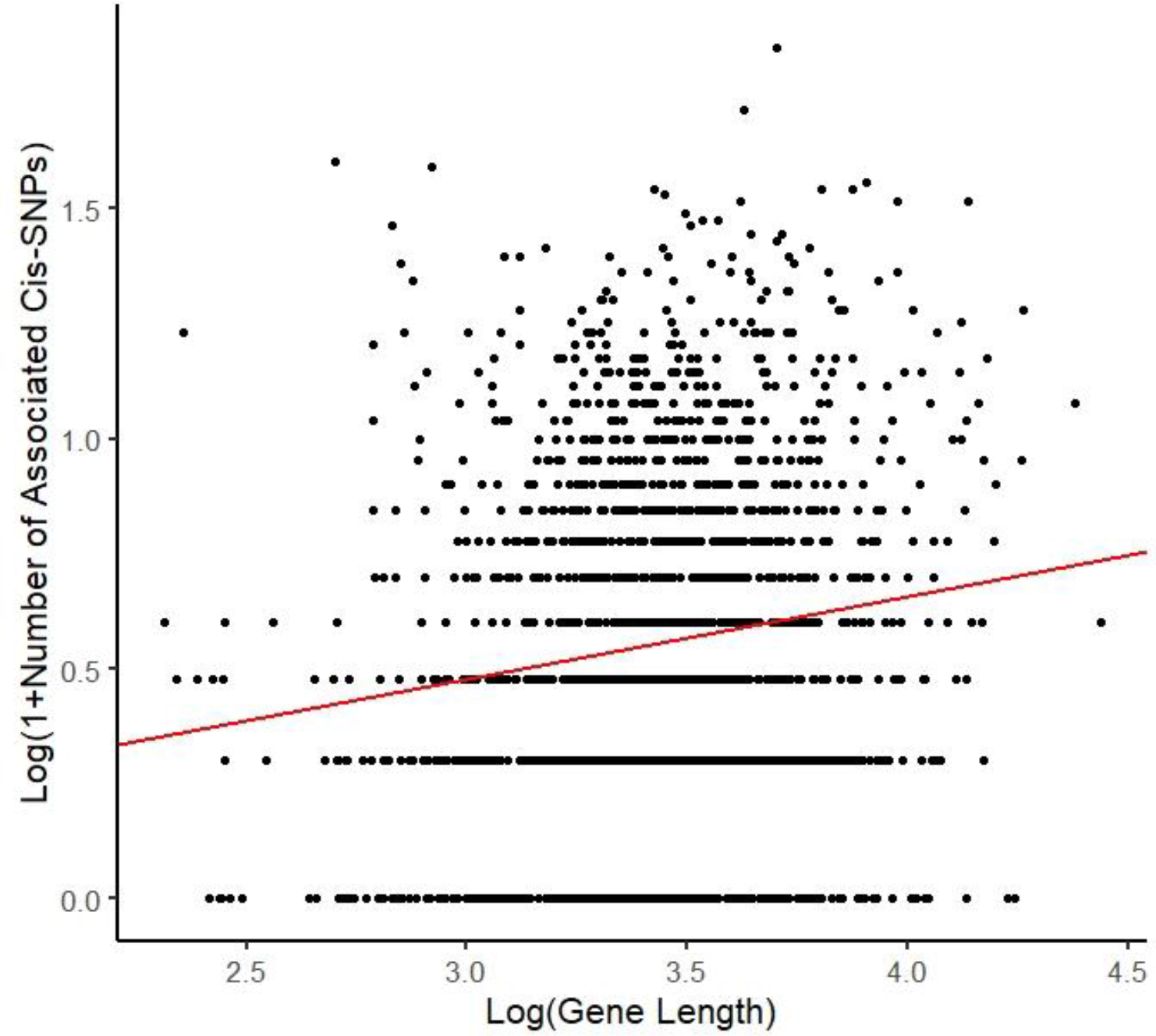
Gene length is positively correlated with the number of SNPs associated with gene expression in cis.

**Figure S9.**
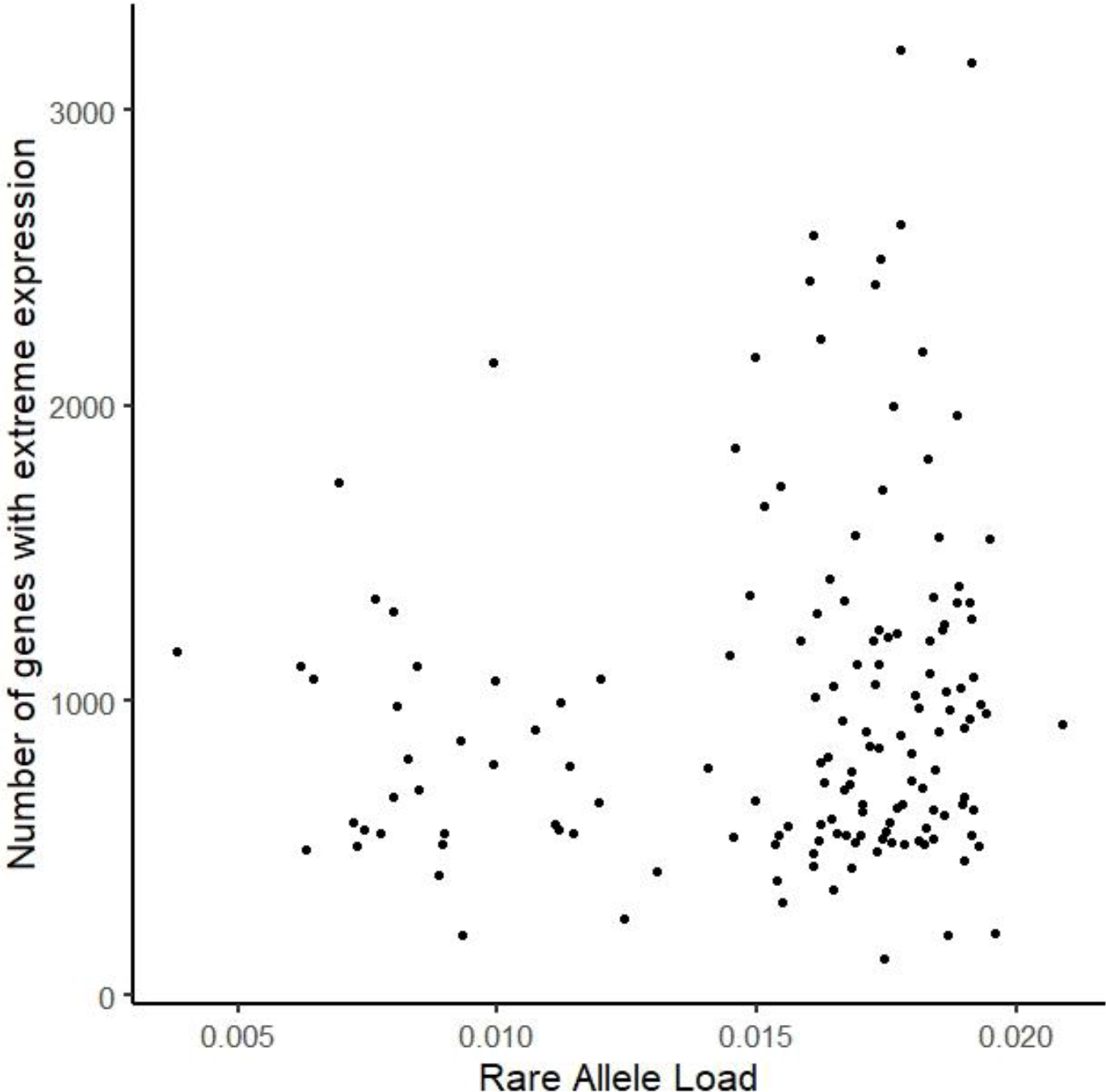
Rare allele load genome-wide does not predict extreme expression transcriptome-wide. For each inbred line, the rare allele load as calculated by Brown and Kelly (2020), is plotted against the number of genes for which a line has extreme expression. Extreme expression describes greater or less than two standard deviations from the mean expression of all lines. We have excluded line Z12, as in the above study, because it is an outlier for rare allele load. Including it does not make the regression significant.

**Figure S10.**
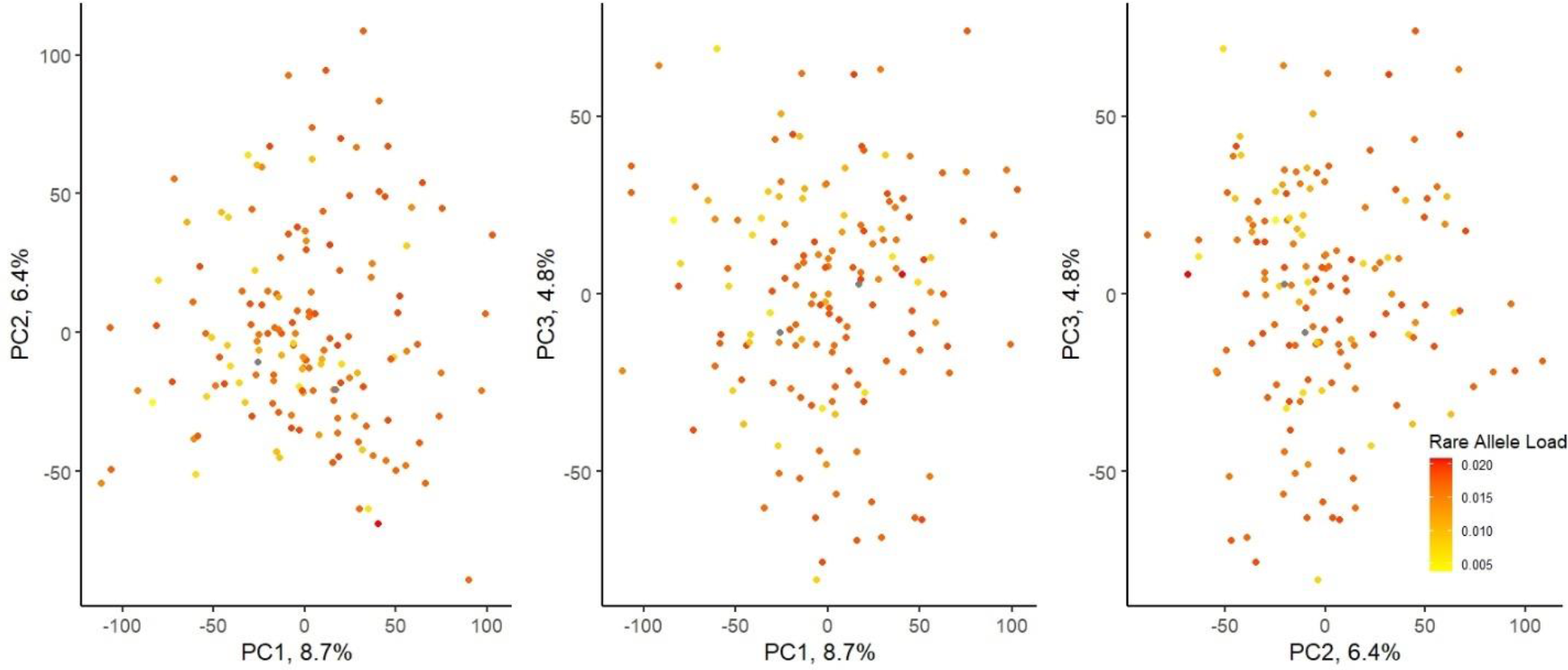
Rare allele load genome-wide does not cluster in gene expression principal component (PC) space. We calculated the first three PCs for gene expression and plot them here, colored by rare allele load of each line. As above, we have excluded line Z12. Variance explained by each PC is labelled on the axes.

**Figure S11.**
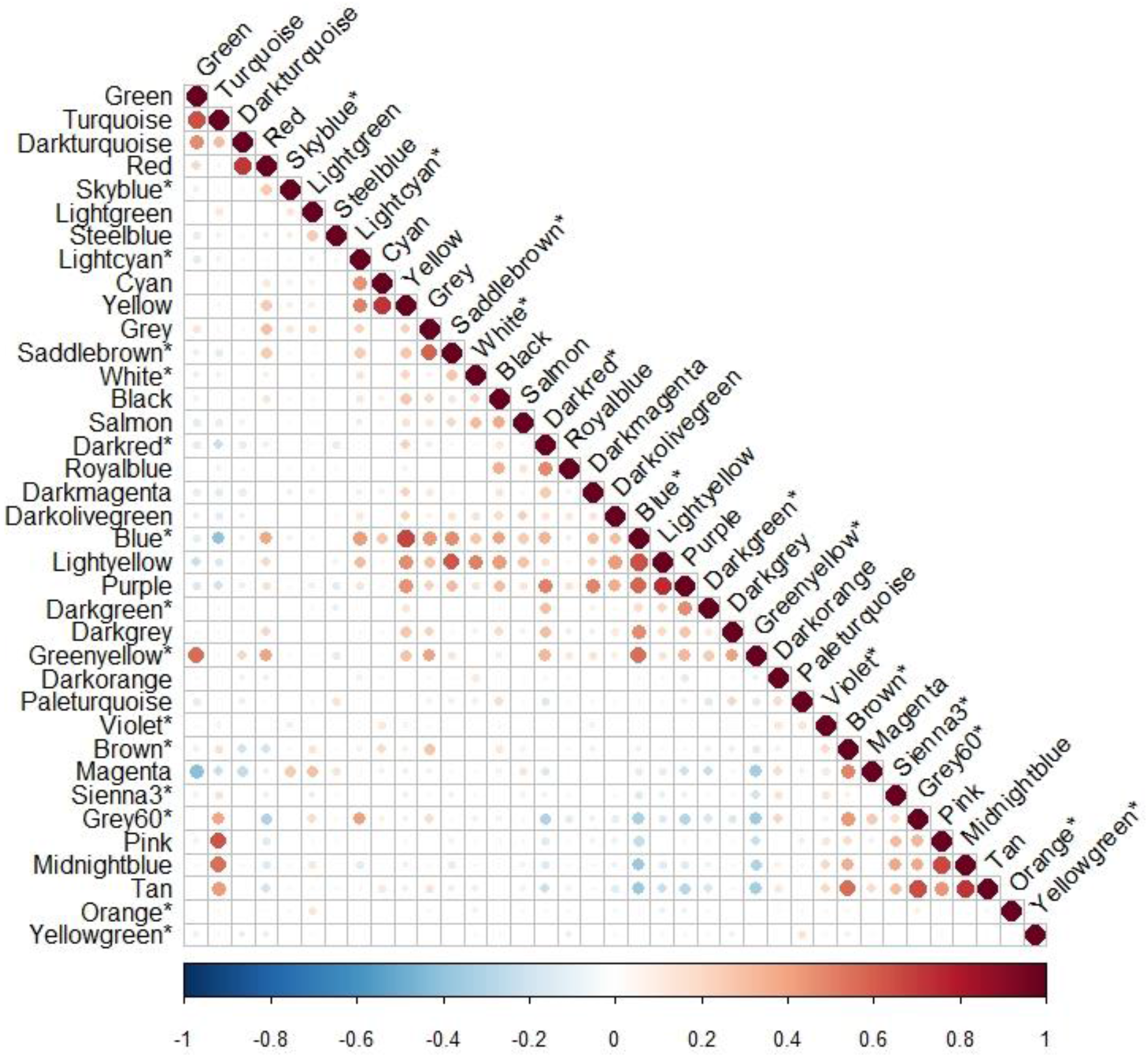
Correlation matrix for gene expression module eigen expression (PC1). A few (20 of 666 comparisons) are correlated at 0.5 < R^2^ < 0.74. Asterisks indicate the 14 modules in common between the modules predicting 4 flower size traits (corolla width and length, anther length, and flower size PC1).

